# TCOF1 is a single-component scaffold of the nucleolar fibrillar center

**DOI:** 10.1101/2022.10.16.512422

**Authors:** Nima Jaberi-Lashkari, Byron Lee, Fardin Aryan, Eliezer Calo

**Affiliations:** Department of Biology, Massachusetts Institute of Technology, Cambridge, United States; David H. Koch Institute for Integrative Cancer Research, Massachusetts Institute of Technology, Cambridge, United States

## Abstract

Many of the biological structures that exist across the tree of life are built on self-interacting scaffolds, from the actin cytoskeleton to the collagen extracellular matrix. Intracellular membraneless organelles, such as the nucleolus, are biological structures consisting of hundreds of dynamically interacting components, yet it is unclear whether the underlying organization of these complex assemblies can be scaffolded by such self-interacting components. Here, we show that TCOF1 is a single-component scaffold of the nucleolar fibrillar center (FC), based on thermodynamics of its assembly in cells, as well as sufficiency and loss-of-function experiments. TCOF1 is necessary for the formation of the FC, and defines the FC through assembly mediated by homotypic interactions of its Serine/Glutamate (S/E)-rich low-complexity regions (LCRs). Ultimately, introduction of TCOF1 into a species that lacks the FC is sufficient to form an FC-like nucleolar subcompartment. Thus, we demonstrate how a single protein component can explain the formation and evolution of a complex biological structure.

## Main Text

The nucleolus is a membraneless organelle containing hundreds of components which are spatially segregated into distinct, nested subcompartments that together carry out sequential steps in the essential process of ribosome biogenesis (*1*). In humans, nucleoli have a tripartite architecture consisting of the fibrillar center (FC), nested within the dense fibrillar component (DFC), which is nested within the granular component (GC). While the GC and DFC are thought to assemble based on many interacting components, including Nucleophosmin (NPM1), Fibrillarin (FBL), and rRNA (*2–4*), the components which underlie the structure of the FC are unknown.

In previous work, we observed that the FC protein TCOF1 was sufficient to assemble and recruit other FC proteins when targeted outside of the nucleolus (*5*). This observation led to the hypothesis that TCOF1 may be capable of self-assembly, and may therefore play a scaffolding role in the FC. Notably, TCOF1 self-assembly would be in contrast to the finding that the highly studied components of intracellular assemblies have been found to not exhibit self-assembly behavior in cells (*6*), leading us to test whether TCOF1 is a self-assembling scaffold using the same thermodynamic framework.

In order to determine whether a given component of an assembly can self-assemble, or if it relies on other components for its assembly, one can measure the thermodynamics of its partitioning in cells (*6*). This can be determined by the relative concentrations of the component in the dense phase (within the assembly) and dilute phase (outside of the assembly), which determine its free energy of transfer (ΔG^transfer^, see Methods), a measure inversely related to the favorability for entering the assembly. Upon increased expression, a component that is dependent on other component(s) in the assembly will reach a point in which it begins to saturate the dense phase and accumulate in the dilute phase. This behavior is referred to as multi-component assembly, and is reflected in an increase in ΔG^transfer^ as well as little to no change in the dense phase size (Fig. 1A). On the other hand, a component capable of self-assembly will partition into the assembly across all expression levels in cells. This behavior is referred to as single-component assembly, in which ΔG^transfer^ remains constant and the dense phase size increases with expression (Fig. 1A).

**Fig. 1.**
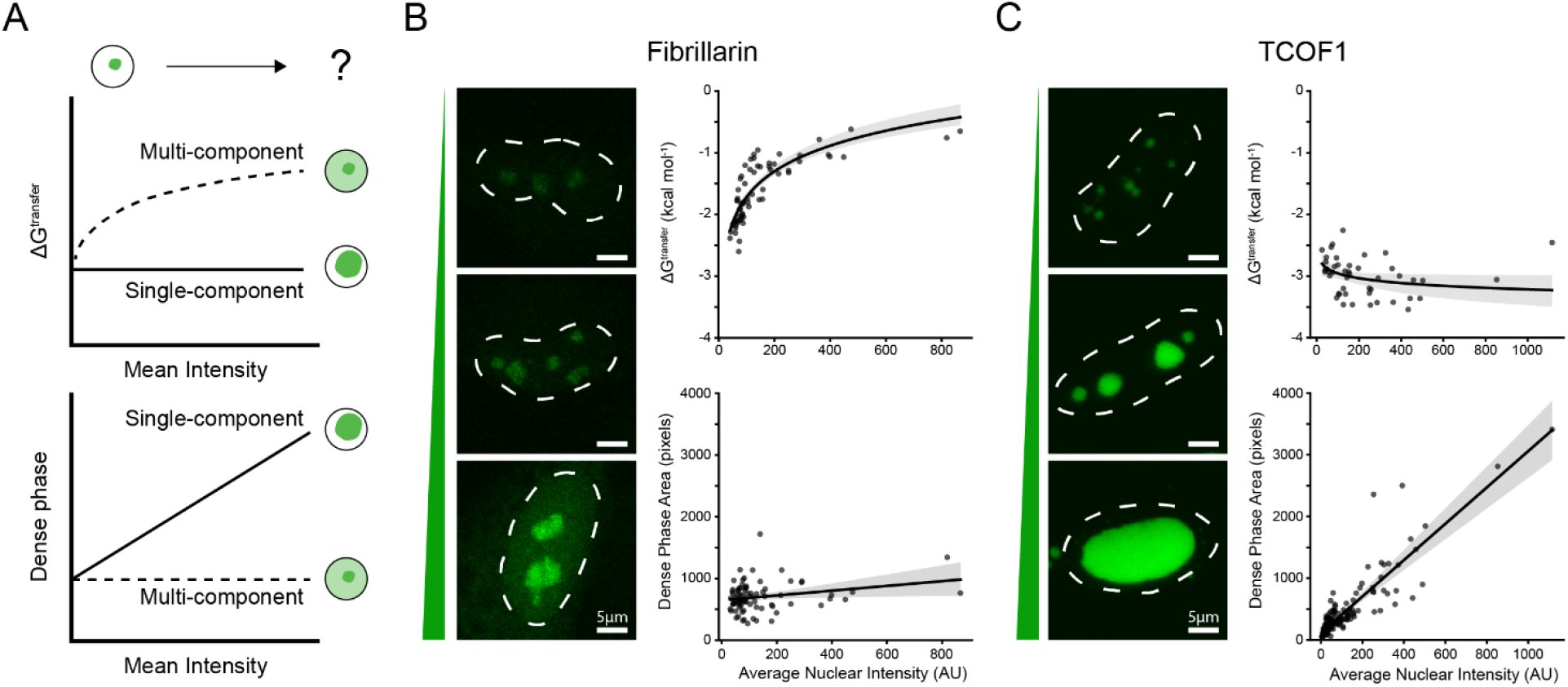
TCOF1 displays single-component assembly in cells. **(A)** Schematic illustrating expected trends of ΔG^transfer^ and dense phase size for proteins with multi-component or single-component assembly behavior. **(B)** Quantification of ΔG^transfer^ and dense phase size vs. average nuclear intensity for meGFP-Fibrillarin. Images of cells with different expression levels are shown on the left. Nuclei are outlined with a dotted line based on Hoechst 33342 signal (not shown). Scale bars are 5 μm. N = 66 nuclei for ΔG^transfer^ measurement, and N = 81 nuclei for dense phase area measurement. Logarithmic and linear fits with corresponding 95% confidence intervals are shown. **(C)** same as (B), but for meGFP-TCOF1. N = 46 nuclei for ΔG^transfer^ measurement, and N = 167 nuclei for dense phase area measurement.

Using this framework, we observed that although Fibrillarin (FBL) and Nucleophosmin (NPM1) displayed multi-component assembly in HeLa cells, TCOF1 displayed single-component assembly. Partitioning of both FBL and NPM1 became unfavorable upon increased expression, as could be seen by increased dilute phase in cells (Fig. 1B, S1A), and both proteins exhibited little to no increase in the size of their dense phases (Fig. 1B, S1A). This observation is consistent with the assembly of FBL and NPM1 depending on other interacting components in nucleoli. On the other hand, TCOF1 maintained its partitioning and increased in dense phase area over roughly 3 orders of magnitude of expression (Fig. 1C). These results argue that TCOF1 is capable of assembling on its own inside cells, representing the first example of a natural protein capable of single-component assembly in an intracellular membraneless organelle.

Strikingly, we observe that components of the fibrillar center maintain their colocalization with TCOF1 assemblies regardless of their size, and simply dilute across larger TCOF1 assemblies (Fig. 2A, Fig. S2A). On the other hand, components of the DFC, GC, and nucleolar rim remain excluded from TCOF1 assemblies (Fig. 2A), suggesting that TCOF1 assemblies maintain an FC identity even as they expand (Fig. 2B). Importantly, another FC protein, RPA43, does not exhibit this behavior (Fig. S2B), which indicates that TCOF1 specifically may underlie the structure of the FC in cells.

**Fig. 2.**
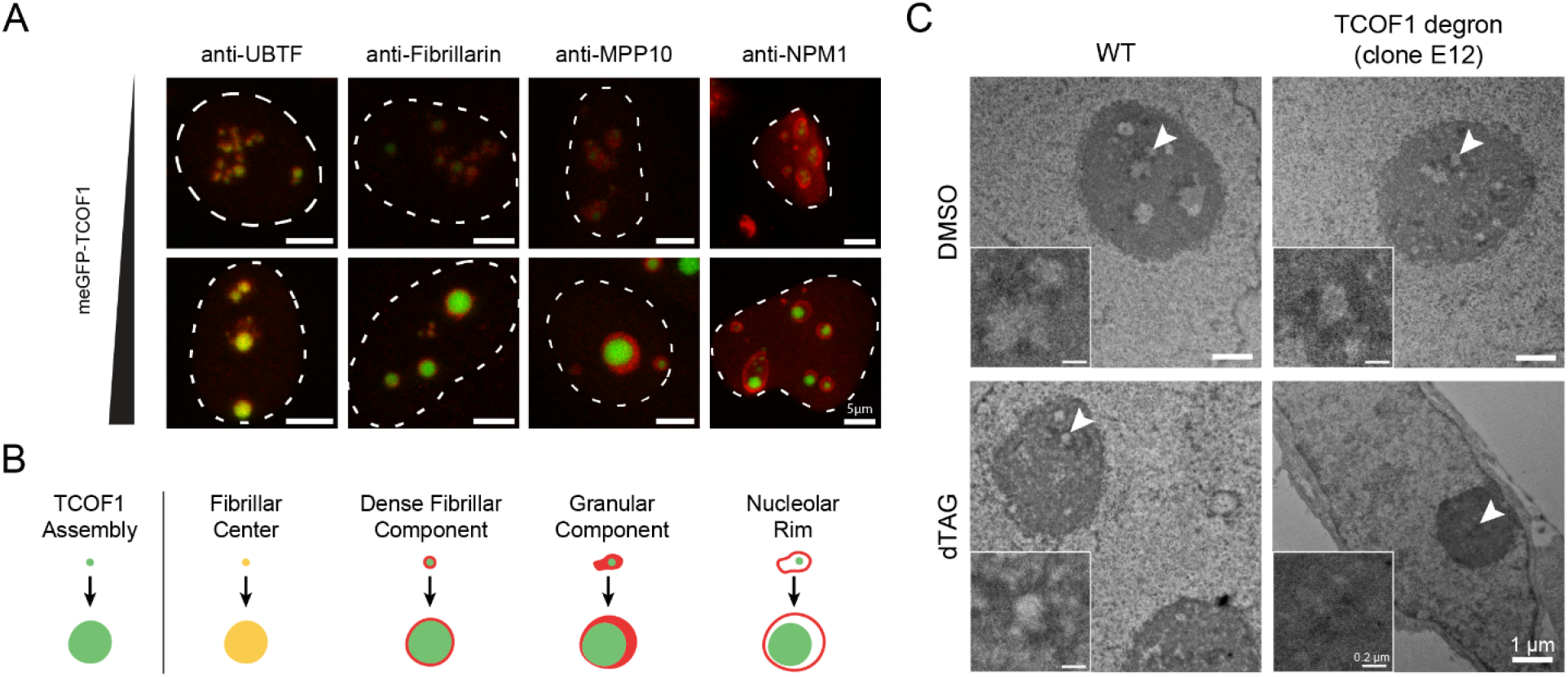
TCOF1 defines the nucleolar fibrillar center. **(A)** Immunofluorescence of endogenous nucleolar subcompartment markers across varying levels of meGFP-TCOF1 expression. Nuclei are outlined with a dotted line based on Hoechst 33342 staining (not shown). Scale bars are 5 μm. **(B)** Model of nucleolar subcompartment structure and FC identity defined by TCOF1 assembly. **(C)** Transmission electron microscopy of WT HeLa cells and TCOF1-FKBP degron HeLa cells in DMSO and dTAG-13 treatment. Scale bar is 1 μm. Inset shows close-up view of region indicated by arrowhead. Inset scale bar is 0.2 μm.

To test the importance of TCOF1 for FC structure, we made a degron-based loss-of-function system for TCOF1 in cells. Specifically, we generated cell lines harboring homozygous in-frame fusions of the FKBP-degron (Fig. S3), which allows for the inducible degradation of degron-tagged proteins (*7*). With the addition of dTAG-13, TCOF1 expression in these lines was reduced to undetectable levels within 4 days (Fig. S4A-C). Moreover, knockdown of TCOF1 reduced levels of nascent nucleolar RNA by 5-ethynyl uridine (5-EU) imaging (Fig. S4D), consistent with previously published studies on the loss of TCOF1 (*8*). The ability to inducibly eliminate TCOF1 allowed us to further assess the role of TCOF1 in FC structure.

The FC, DFC, and GC are defined by their unique ultrastructural appearance under transmission electron microscopy (Fig. S5A). We observed through electron microscopy (EM) that the addition of dTAG-13 in two separate TCOF1 degron lines resulted in ablation of the FC, while having no effect on WT HeLa cells, which do not contain the FKBP degron (Fig. 2C, S5B). Notably, nucleoli in dTAG-treated TCOF1 degron cell lines still contained both the GC and DFC comparable to the WT HeLa cells and DMSO-treated TCOF1 degron cells (Fig. 2C, S5B), showing that TCOF1 is specifically required for the FC.

The observations above suggest that the assembly of this single protein is tied to the assembly of the FC as a whole, such that it defines the FC. Thus, by understanding the molecular mechanism of TCOF1 assembly, we can understand the molecular structure of the FC. We therefore set out to determine the features of TCOF1 which allow it to assemble in cells.

Given the importance of valency for the formation of biological assemblies (*9*), we hypothesized that the central repeat domain of TCOF1 (Fig. S6A) may be involved in its assembly. A version of TCOF1 with this region deleted, TCOF1 Δ83-1170, failed to assemble in a single-component manner (Fig. 3A), showing that the repeat domain is essential for the single-component assembly of TCOF1.

**Fig. 3.**
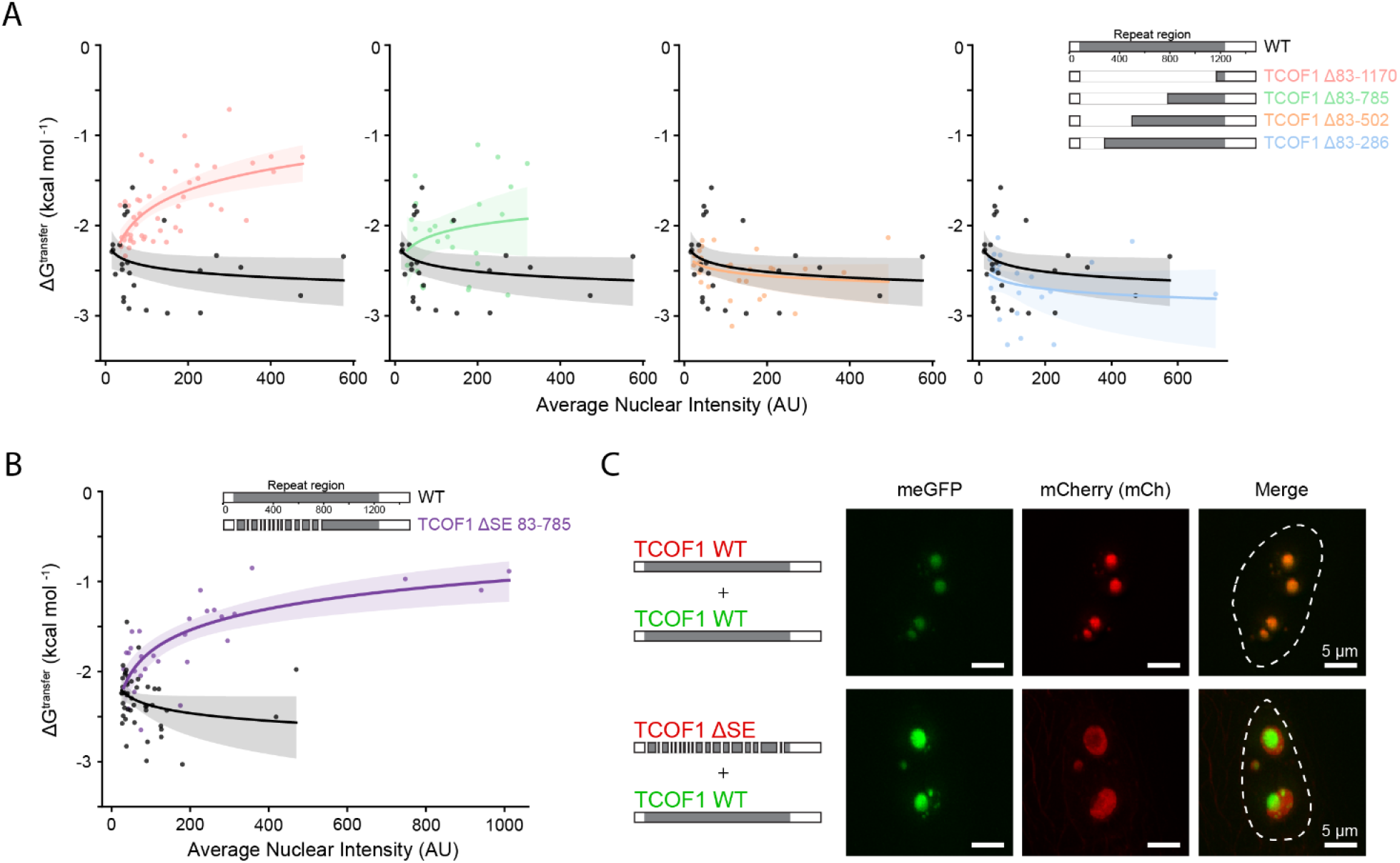
TCOF1 self-assembly is mediated by self-interacting S/E-rich LCRs. **(A)** Quantification of ΔG^transfer^ vs. average nuclear intensity for WT TCOF1 and TCOF1 mutants with internal deletions in the central repeat region. Data for WT TCOF1 is shown on each plot for clarity. N = 26 nuclei for WT TCOF1, N = 48 nuclei for TCOF1 Δ83-1170, N = 25 nuclei for TCOF1 Δ83-785, N = 27 nuclei for TCOF1 Δ83-502, N = 24 nuclei for TCOF1 Δ83-286. Logarithmic fits and corresponding 95% confidence intervals are shown. **(B)** Quantification of ΔG^transfer^ vs. average nuclear intensity for WT TCOF1 and TCOF1 ΔSE 83-785. N = 46 for WT TCOF1 and N = 29 for TCOF1 ΔSE 83-785. **(C)** Fluorescence microscopy of co-assembly experiments in which the indicated mCherry and meGFP constructs were co-transfected into HeLa cells. Nuclei are outlined with a dotted line based on Hoechst 33342 signal (not shown). Scale bars are 5 μm.

Serial truncations of this domain resulted in eventual loss of single-component assembly (Fig. 3A, S6B, S7A). This could not be explained by the direction from which the truncations were made (Fig. 3A, S6B, S7A), or by any single part of the repeat domain shared between constructs capable of single-component assembly (Fig. S6C). Given that the region necessary for single-component assembly could not be mapped to a single location within this repeat domain (Fig. S6D), we reasoned that it is likely dispersed along the sequence.

The defining feature of the central repeat domain of TCOF1 is the existence of 16 roughly evenly spaced S/E-rich low complexity regions (LCRs) (Fig. S6A). We hypothesized that these LCRs are the dispersed feature necessary for TCOF1’s single-component assembly behavior. Supporting this hypothesis, we observed a strong relationship between the number of S/E-rich LCRs present in each of our mutants and whether or not the mutants assembled in a single- or multi-component matter (Fig. S6D, S7A). Mutants with 9 or more copies of S/E-rich LCRs maintained single-component assembly indistinguishable from WT TCOF1, while those with 5 or fewer copies of the S/E rich LCRs failed to assemble in a single-component manner (Fig. S6, S7A) (*10*). To explicitly test the role of the S/E-rich LCRs in the assembly of TCOF1, we generated a TCOF1 mutant in which all but 5 S/E-rich LCRs were specifically deleted (TCOF1 ΔSE 83-785). We observed that TCOF1 ΔSE 83-785 failed to assemble in a single-component manner (Fig. 3B, S6D), showing that this assembly behavior of TCOF1 requires its S/E-rich LCRs.

Moreover, when assessing the ability of the truncation mutants to expand the fibrillar center, mutants with more S/E-rich LCRs appeared to exclude the GC (as assessed by MPP10), while those with fewer S/E-rich LCRs were unable to do so (Fig. S7B), suggesting that the ability of TCOF1 to assemble is intertwined with its ability to define the FC. If so, then specific perturbations to TCOF1 assembly by deletion of the S/E-rich LCRs would be expected to also disrupt its ability to define the FC. Indeed, a mutant in which all of the 16 S/E-rich LCRs were deleted (TCOF1 ΔSE) failed to exclude the GC marker MPP10 (*11*), similar to the mutants of TCOF1 which failed to assemble (Fig. S7B). This result not only shows that the S/E-rich LCRs are necessary for TCOF1 to define the FC, but also suggests that TCOF1 assembly is inherently tied to the formation of the FC as a compartment.

The coupling between TCOF1 assembly and the formation of the FC raises the question of how S/E-rich LCRs facilitate TCOF1 assembly. We reasoned that if TCOF1 self-assembles via homotypic interactions between its S/E-rich LCRs, then two versions of TCOF1 will not be able to co-assemble in cells unless both of them possess S/E-rich LCRs. This is because interactions between the two versions will not be possible when either lacks all of its S/E-rich LCRs. However, if TCOF1 self-assembles via heterotypic interactions between S/E-rich LCRs and other regions in TCOF1, then a version of TCOF1 which lacks the S/E-rich LCRs will still co-assemble with a version of TCOF1 which contains the S/E-interacting element, as interactions will still be possible between these two versions.

To determine if S/E-rich LCRs mediate assembly by homotypic or heterotypic interactions within TCOF1, we co-expressed meGFP- and mCherry-tagged versions of TCOF1 to assess their co-assembly. When co-expressed in cells, WT TCOF1 can co-assemble with itself, as seen by the colocalization of meGFP and mCherry signal (Fig. 3C, top row). However, when we co-expressed WT TCOF1 with TCOF1 ΔSE, they failed to co-assemble. Rather, WT TCOF1 was able to exclude TCOF1 ΔSE (Fig. 3C, bottom row), suggesting that TCOF1’s S/E-rich LCRs mediate assembly through self-interaction, and not by interacting with another element within TCOF1.

Collectively, these results provide a molecular mechanism by which TCOF1 defines the nucleolar FC, through self-assembly mediated by homotypic interactions of the repeated S/E-rich LCRs. These findings show that the size, shape, and even presence of the FC in cells can be influenced through the precise expression, or lack of expression, of TCOF1. This structural role of TCOF1 in the FC led us to ask whether our understanding of TCOF1 can provide a molecular explanation for the evolutionary origins of the FC.

Many eukaryotes possess bipartite nucleoli, which lack an FC-like subcompartment (*12, 13*). While tripartite nucleoli contain three subcompartments (an FC, DFC, and GC), bipartite nucleoli contain only GC-like and DFC-like subcompartments referred to as Granular and Fibrillar Zones (GZ and FZ), respectively. Although these structural similarities have been observed between tripartite and bipartite nucleoli, the underlying differences between these nucleolar organizations has been a longstanding question, subject to much speculation (*13*). Given that the FC is thought to have emerged only in amniotes (*12*), we hypothesized that TCOF1 may have emerged at a similar point in evolution.

We first broadly assessed whether the presence of TCOF1 orthologs correlates with the presence of a nucleolar fibrillar center (i.e. tripartite nucleolus). To do this, we looked for species for which we could assess both nucleolar structure by existing transmission electron microscopy data, and presence or absence of TCOF1 orthologs in the proteome (see Methods). Of the eight species we looked at, we observe a relationship between the existence of an FC in nucleoli and the presence of TCOF1 orthologs (Fig. 4A, Fig. S9B-F, Table S1, Methods), with most species we looked at either having both a TCOF1 ortholog and an FC, or lacking both (Fig. 4A).

**Fig. 4.**
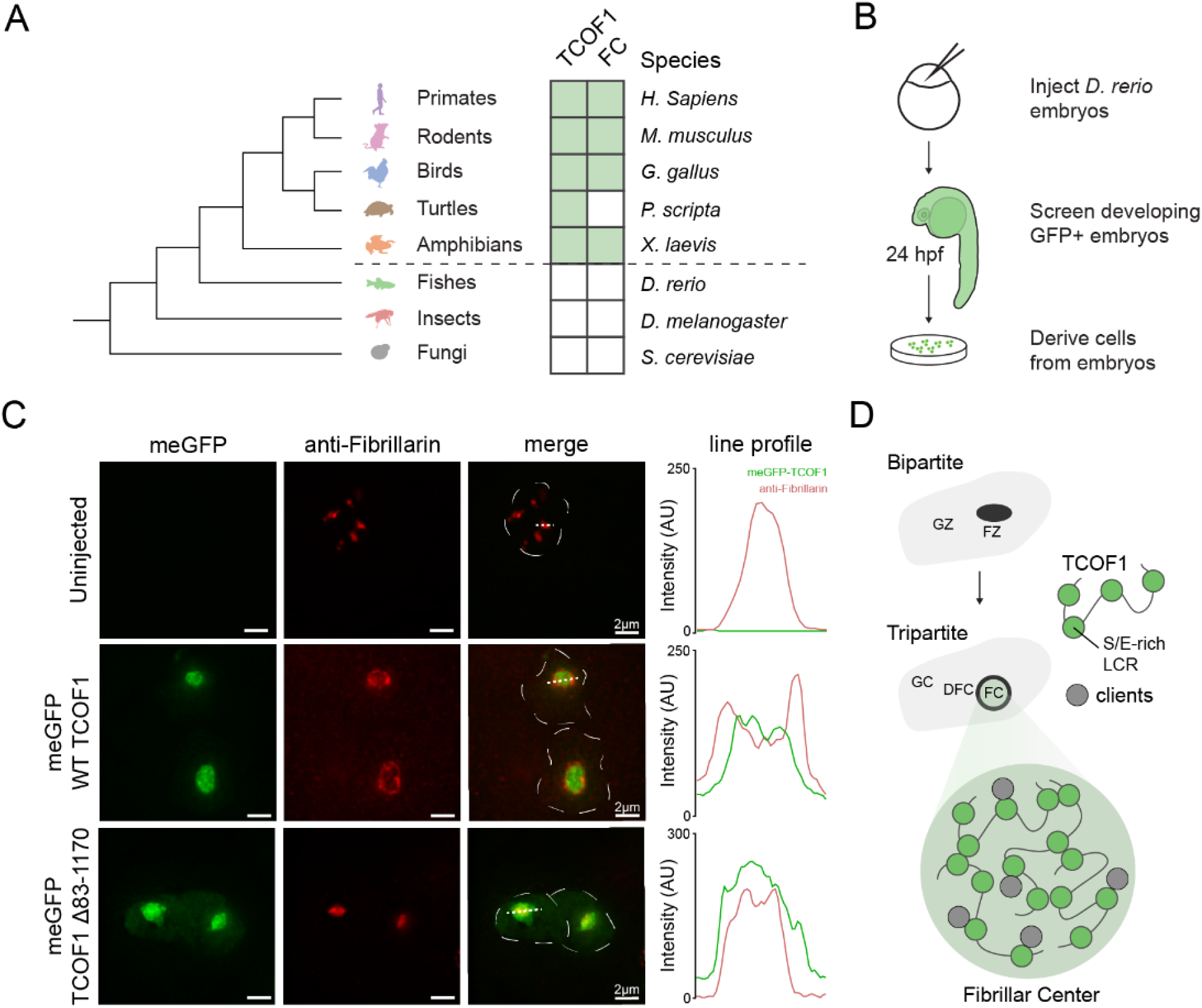
Introduction of human TCOF1 into zebrafish embryos results in the formation of an FC-like compartment. **(A)** Analysis of presence of TCOF1 and tripartite (FC-containing) nucleoli in species as indicated. Tree on the left illustrates the relative evolutionary relationships between species. **(B)** Schematic of approach to assess TCOF1 assembly in cells derived from zebrafish embryos. See Materials and Methods for details. **(C)** Fluorescence microscopy of cells derived from uninjected zebrafish embryos, or zebrafish embryos injected with meGFP-TCOF1 mRNA or meGFP-TCOF1 Δ83-1170 mRNA. Fibrillar zone is visualized using Fibrillarin antibody. Nuclei are outlined with a dotted line based on Hoechst 33342 signal (not shown). Scale bars are 2 μm. Line profile plots are included over the region indicated by the finely dotted line for meGFP (green) and Fibrillarin channels (red). D) Model for the emergence and underlying structure of the FC.

Given that the emergence of the FC is thought to have occurred in amniotes, we focused on species which span this transition, the bony fish D. rerio, the amphibian X. laevis, the turtle P. scripta, and the bird G. gallus (Fig. 4A). Consistent with the hypothesis of co-emergence of FC and TCOF1, D. rerio lacks both TCOF1 and an FC, while G. gallus has both. Although P. scripta possesses TCOF1 but lacks an FC, the existence of a TCOF1 ortholog and the ability to form an FC in X. laevis (*14*) suggests that this may be due to a loss of FC in turtles for other reasons. This association between the presence of TCOF1 and the FC led us to the hypothesis that emergence of TCOF1 may have played a role in the emergence of the FC across evolution. If true, this hypothesis would suggest that introduction of TCOF1 into a bipartite system may be sufficient for the formation of an FC-like nucleolar compartment.

To test this, we asked whether introduction of human TCOF1 into zebrafish, which lack both TCOF1 and an FC (Fig. 4A, Fig. S9A), could result in the formation of an FC-like compartment. We injected mRNA encoding meGFP-TCOF1 into one-cell stage zebrafish embryos, and 24 hours after fertilization we derived cells from these embryos and performed immunofluorescence of the DFC/FZ marker Fibrillarin (Fig. 4B). In uninjected embryos, Fibrillarin staining appears as small nuclear bodies, consistent with its localization to the FZ of bipartite nucleoli. Strikingly, in cells from embryos injected with WT TCOF1 mRNA, we observed the formation of nuclear TCOF1 assemblies, nested within Fibrillarin (Fig. 4C). In these cells, Fibrillarin organization conforms to the shape of the TCOF1 assembly, surrounding it as a DFC would surround an FC in tripartite nucleoli. In fact, we see that Fibrillarin exclusion is maintained as these TCOF1 assemblies expand, akin to the expanded FCs we observed in human cells, which have tripartite nucleoli (Fig. 2A, S10B). Importantly, introduction of an assembly-deficient mutant of TCOF1 lacking the central repeat domain into zebrafish embryos resulted in mixing of mutant TCOF1 with Fibrillarin (Fig. 4C) suggesting that the ability of human TCOF1 to form an FC-like compartment in zebrafish is dependent on its ability to self-assemble. The discovery that a protein component, TCOF1, is sufficient to form a FC-like compartment in bipartite nucleoli is to our knowledge the only direct experimental support for a molecular explanation of the bipartite to tripartite transition, through the emergence of TCOF1.

Together, this work presents a model for the structure and molecular origins of the nucleolar FC, in which emergence of TCOF1 is coupled to the emergence of the FC, the underlying structure of which is defined by the ability of TCOF1 to self-assemble through homotypic interactions between its S/E-rich LCRs (Fig. 4D). Future studies will leverage this understanding to gain insight into FC function. More generally, this work not only suggests that a complex biological structure may be formed by the ability of a protein to act as a single-component scaffold, but also presents a general model for the formation and origin of these complex structures.

The existence of TCOF1 as a single-component scaffold in cells suggests that membraneless organelles need not always form from a large number of co-interacting molecules, raising the question of if other membraneless organelles are truly scaffolded by such interactions, or if more thorough screening is needed to determine their underlying single-component scaffolds. The idea that complex assemblies within the cell may be scaffolded by a single protein is consistent with an emerging view that the underlying principles of intracellular membraneless compartments may relate to those in other biomolecular structures (*5*), such as the siliceous skeletons of certain sponges (*15*) or spider silk. These assemblies often form from very few scaffolding components, allowing the properties of the assembly to be largely controlled by the expression of just a few components. For example, while spider silk does contain other components which influence the specific physical properties of the silk fibers (*16*), the formation of this assembly is dependent on the expression of two spidroin proteins (*17–19*), which are sufficient to form silk fibers (*20, 21*). Likewise, we find that the size of the nucleolar fibrillar center appears to be tuned by the expression level of TCOF1 protein. These findings raise the possibility that other intracellular membraneless organelles may be similarly tuned and organized.

Given that emergence of an assembly based on single-component scaffolds requires only a single self-assembling component, these assemblies may arise in fewer evolutionary steps when compared to ones with a large number of co-interacting molecules. Following emergence of a single-component scaffold, molecules which contribute the structure or function of the assembly can be gained in a stepwise manner over time. In fact, assemblies thought to be made up of highly interconnected components, such as stress granules, p-bodies, and the nucleolar GC (*3, 6, 22, 23*), are much more evolutionarily ancient than the FC, which relies on TCOF1. Similarly, several species specific (and therefore more recently emerged) assemblies, such as spider silk, are scaffolded by just a few components (*17*). These distinct assembly architectures may potentially be explained by a more general model for the emergence of complex biological assemblies, in which assemblies first arise from single self-interacting components, gain interaction partners over time, and eventually rely on these interaction partners for the structure of the assembly itself.

While we are only beginning to gain insight into these diverse complex biological assemblies, our understanding of TCOF1 as an archetypal single-component scaffold may inform future studies of the structure and function of biological assemblies across the tree of life.

## Acknowledgments

We would like to thank all members of the Calo lab for helpful discussions and feedback on the manuscript. We would also like to thank the Swanson Biotechnology Center Microscopy and Flow Cytometry Facilities, as well as the Harvard Medical School EM Facility. We thank the Burge and Boyer labs at MIT for providing FKBP degron plasmids and dTAG-13, respectively.

## Author contributions

Conceptualization: NJ, BL; Data Curation: NJ, BL; Formal Analysis: NJ, BL; Funding acquisition: EC; Investigation: NJ, BL, FA; Methodology: NJ, BL, FA; Resources: EC; Software: NJ, BL; Supervision: NJ, BL; Validation: NJ, BL; Visualization: NJ, BL; Writing – original draft: NJ, BL, FA; Writing – review & editing: NJ, BL, EC;

## Data and materials availability

All data and materials are available upon request for the purposes of reproducing or extending the analysis. Code used for image analysis can be found on zenodo at: https://doi.org/10.5281/zenodo.7236056

## Materials and Methods

### Cell lines

Hela cells were obtained from ATCC. Cells tested negative for mycoplasma.

### Plasmids and constructs

All constructs contain a GSAAGGSG peptide linker between GFP and the protein of interest. Coding sequences of wildtype proteins were cloned from human cDNA, from which deletions were made. The only exception was pcDNA3.1(+) meGFP - TCOF1 ΔSE and ΔSE 83-785, which were cloned from a synthesized gene block (Twist Bioscience) which was codon optimized to reduce repetitiveness of the DNA sequence.

| Plasmid | Source | Identifier |
| --- | --- | --- |
| pcDNA3.1(+) eGFP - NPM | This paper | RP14 (NPM1 WT) |
| pcDNA3.1(+) meGFP - FBRL | This paper | RP162 (FBRL WT) |
| pcDNA3.1(+) meGFP - RPA43 | Lee, Jaber-Lashkari et al. 2022 | RP104 (RPA43 WT) |
| pcDNA3.1(+) meGFP - TCOF1 | Lee, Jaber-Lashkari et al. 2022 | RP133 (TCOF1 WT) |
| pcDNA3.1(+) meGFP - TCOF1 ΔSE (ΔS83-E97, S171-S178, D197-D211, E267-E282, E336-E352, E407-E422, E472-E487, E542-E559, S608-E622, E691-E704, E760-E773, E865-E882, D930-E944, S990-D1005, E1141-E1161, E1185-D1195) | This paper | RP136 (TCOF1 ΔSE) |
| pcDNA3.1(+) meGFP - TCOF1 (ΔS83-G286) | This paper | RP165 (TCOF1 ΔS83-G286) |
| pcDNA3.1(+) meGFP - TCOF1 (ΔS83-K502) | This paper | RP166 (TCOF1 ΔS83-K502) |
| pcDNA3.1(+) meGFP - TCOF1 (ΔS83-V785) | This paper | RP167 (TCOF1 ΔS83-V785) |
| pcDNA3.1(+) meGFP - TCOF1 (ΔS83-H1170) | This paper | RP168 (TCOF1 ΔS83-H1170) |
| pcDNA3.1(+) meGFP - TCOF1 (ΔR688-S1234) | This paper | RP170 (TCOF1 ΔR688-S1234) |
| pcDNA3.1(+) meGFP - TCOF1 (ΔR405-S1234) | This paper | RP171 (TCOF1 ΔR405-S1234) |
| pcDNA3.1(+) meGFP - TCOF1 (ΔT167-S1234) | This paper | RP172 (TCOF1 ΔT167-S1234) |
| pcDNA3.1(+) meGFP - TCOF1 (ΔS503-Q687) | This paper | RP177 (TCOF1 ΔS503-Q687) |
| pcDNA3.1(+) meGFP - TCOF1 ΔSE 83-785 (ΔS83-E97, S171-S178, D197-D211, E267-E282, E336-E352, E407-E422, E472-E487, E542-E559, S608-E622, E691-E704, E760-E773) | This paper | RP192 (TCOF1 ΔSE 83-785) |
| pcDNA3.1(+) mCherry - TCOF1 | This paper | RP197 (TCOF WT) |
| pcDNA3.1(+) mCherry - TCOF1 ΔSE<br>(ΔS83-E97, S171-S178, D197-D211,<br>E267-E282, E336-E352, E407-E422,<br>E472-E487, E542-E559, S608-E622,<br>E691-E704, E760-E773, E865-E882,<br>D930-E944, S990-D1005, E1141-E1161,<br>E1185-D1195) | This paper | RP198 (TCOF ΔSE) |

### Cell culture

HeLa cells were cultured in 5% CO2 on cell culture-treated 10 cm plates (Genesee Scientific, 25-202) in Dulbecco’s Modified Eagle Medium (DMEM, Genesee Scientific, 25-500) supplemented with 10% Fetal bovine serum (FBS, Gemini Bio-products, 100-106) and 1% Penicillin/Streptomycin (Gibco, 10378-016). Cells were split 1:10 every 3 days by using trypsin (Gibco, 25200072).

### Generation of degron cell lines

Degron lines were generated in HeLa cells using sequential rounds of the following approach of CRISPR-Cas9-mediated cutting, homologous recombination (HR)-mediated tagging with the FKBP degron (*7*) fused to different selection markers, and selection, to achieve complete tagging of the TCOF1 locus. See Figure S3A for flowchart, and Figure S3B for HR templates.

An sgRNA targeting the 3’UTR of TCOF1 proximal to the stop codon was cloned into the Cas9-containing plasmid PX458 (Addgene, 48138) by golden gate cloning, and subsequently sequence verified. The sgRNA used (5’-GTATGACGAGCACCAGCACC-3’) had on-target and off-target scores of 58.8 and 57.8, respectively, and was designed using software on Benchling. Off-target scores calculation included masked and low-complexity genomic regions.

HR templates depicted in Figure S3B are described here for completeness. HR templates contained an upstream homology arm 900 bp in length, which corresponds to the 900 bp of the TCOF1 genomic locus immediately upstream of the stop codon, and was cloned from HeLa genomic DNA. Downstream of this homology arm, we cloned the FKBP degron cassette consisting of (GSG linker)-(FKBP-V)-(2xHA)-(GSG linker)-(P2A)-(Selection marker)-(STOP). The stop codon was followed by a downstream homology arm, which corresponded to the 821 bp of the TCOF1 genomic locus immediately downstream of the stop codon, and was cloned from HeLa genomic DNA. In the downstream homology arm, the PAM site at which the selected sgRNA cuts was mutated from AGG to AGC to prevent cutting of the locus after successful tagging. The entire HR template was cloned into a plasmid.

The process for generating the lines is depicted in Figure S3A, and described here. First, HeLa cells were co-transfected with the TCOF1 sgRNA plasmid and the HR template carrying Hygromycin resistance. Two days after transfection, cells were selected with 375 μg/mL of Hygromycin for 1 week, generating a Hygromycin resistant (HygR) population. This HygR population was then used in two independent tagging processes, with either Blasticidin- or Puromycin-containing versions of the HR-template, which led to the generation of the E6 and E12 clones used in the study.

For generation of the E6 clone, the HygR population was co-transfected with the TCOF1 sgRNA population and the HR template carrying Blasticidin resistance. Two days after transfection, cells were selected with 4 μg/mL of Blasticidin, for 15 days, generating a HygR, BlastR resistant population. This HygR BlastR population was then single-cell sorted into 96-well plates using a BD FACSAria cell sorter and grown up in the absence of selection. Clones were grown up and screened by PCR, followed by western blotting and immunofluorescence, as described in ‘Validation of degron cell lines’ below, and Clone E6 was selected for use in subsequent experiments.

For generation of the E12 clone, the HygR population was co-transfected with the TCOF1 sgRNA population and the HR template carrying Puromycin resistance. Two days after transfection, cells were selected with 2 μg/mL of Puromycin, for 15 days, generating a HygR, PuroR resistant population. This HygR PuroR population was then single-cell sorted into 96-well plates using a BD FACSAria cell sorter and grown up in the absence of selection. Clones were grown up and screened by PCR, followed by western blotting and immunofluorescence, as described in ‘Validation of degron cell lines’ below, and Clone E12 was selected for use in subsequent experiments.

### Validation of degron cell lines

Degron cell lines were first screened by PCR, and then further validated by western blot and immunofluorescence (Fig. S3, S4). Putative clones, including clones E6 and E12, were identified in the PCR screening, looking for lack of a wildtype PCR product (1268 bp) using genotyping primers spanning the genomic locus outside of the left homology arm and inside the right homology arm (forward: 5’-GCTGGCCTCCAGGGGGCAGGTGAA-3’; reverse primer: 5’-ACAGGGGACACCAGAGCTGT-3’). Following PCR screening, clones were tested for TCOF1 degradation upon dTAG-13 treatment (24hrs, 500 nM) by western blot as well as immunofluorescence. Two independent clones, E6 and E12, were selected on the basis of 1) a lack of a wild-type band by PCR screening, 2) loss of TCOF1 protein after dTAG treatment, and 3) nucleolar localization which is lost after dTAG treatment by immunofluorescence.

### Degron knockdown and dTAG-13 treatment

In experiments using dTAG-13 treatment to induce knockdown, 24 hours after plating, cells were treated with DMSO or 500 nM dTAG-13 for 4 days, replacing media once 2 days after the initial treatment. For these experiments, cells were plated at low seed densities to account for the 4 days of treatment. For immunofluorescence assays in 24-well plates, cells were seeded at 12,500 cells per well. For western blots in 6-well plates, cells were seeded at 75,000 cells per well.

### Western blotting

Cells were collected for whole cell protein lysate using RIPA buffer (150 mM NaCl, 5 mM EDTA, 50 mM TRIS pH 8.0, 0.5% Sodium deoxycholate, 1% NP-40, 0.1% SDS) with 1x cOmplete protease inhibitor cocktail (Sigma Aldrich, 11697498001) and 1mM PMSF (ThermoFisher Scientific, 36978). Lysate was sonicated using a BioRuptor 300 for 5 minutes (5 cycles of 30 secs on, 30 secs off) and cleared by taking the supernatant after spinning at 15,000 x g. Total protein concentrations were measured by Bradford Assay (Thermo Fisher Scientific, 23246), and 50 μg of total protein was run on a Tris-glycine polyacrylamide gel. Transfer to a methanol-activated PVDF membrane was run in Tris-glycine buffer (1x Tris glycine, 10% methanol, 0.05% SDS) at 40 V for 16 hours at 4°C. After transfer, membranes were blocked in 5% milk in PBST (1x PBS, 0.1% Tween-20) for 1 hour and blotted with primary antibody (anti-TCOF1, Proteintech, 11003-1-AP; anti-HA, Cell Signaling (C29F4), 3724S; anti-Tubulin, Thermo Scientific, MA5-16308) overnight at 4°C. Membranes were then blotted with secondary antibodies (anti-rabbit, Invitrogen, 32260; anti-mouse, Invitrogen, 32230) for 1 hour at room temperature, developed using SuperSignal West Femto (Thermo Scientific, 34096), and imaged on a Bioanalytical Imaging System model c500 (Azure Biosciences).

### Fluorescence microscopy

Glass coverslips (Fisherbrand, 12-545-80) were placed in 24-well plates (Genesee Scientific, 25-107) and coated in 3 μg/mL of fibronectin (EMD Millipore, FC010) for 30 minutes at room temperature. HeLa cells were seeded in each well at 50,000 cells per well. 24 hours after seeding, the cells were transfected with GFP-tagged protein plasmids using Lipofectamine 2000 (Invitrogen, 11668027). Each well was transfected using 100 ng of plasmid and 1 μL of Lipofectamine 2000 in a total of 50 μL of OptiMEM (Gibco, 31985070) according to the Lipofectamine 2000 instructions. For co-assembly experiments, 100 ng of each plasmid (200 ng total) was transfected using the same mix.

Cells on glass coverslips were collected for immunofluorescence 48 hours after transfection. Cells were collected by washing with 1x PBS (Genesee Scientific, 25-508) and fixation in 4% paraformaldehyde (PFA) for 15 minutes at room temperature, followed by another 3 washes with 1x PBS.

In experiments with costaining for markers by immunofluorescence, cells were permeabilized and blocked by incubation in blocking buffer (1% BSA (w/v), 0.1% Triton X-100 (v/v), 1x PBS) for 1 hour at room temperature. Coverslips were then incubated overnight at 4°C in a 1:100 dilution of primary antibody (anti-UBTF, Novus Biologicals, NBP1-82545; anti-POLR1A, Novus Biologicals, NBP2-56122; anti-Fibrillarin, EMD Millipore, MABE1154; anti-MPP10, Novus Biologicals, NBP1-84341; anti-Nucleophosmin, Abcam, ab86712; anti-TCOF1, Novus Biologicals, NBP1-86909; anti-HA, Invitrogen, 26183) in blocking buffer. After 3 washes with blocking buffer, coverslips were incubated for 2 hours in a 1:1000 dilution of secondary antibody (anti-rabbit Alexa 647, Invitrogen, A-27040; anti-mouse Alexa 647, Invitrogen, A-32728, anti-mouse Alexa 488, Invitrogen, A-32723). Coverslips were washed 3 times with blocking buffer, and then once with 1x PBS.

Nuclei were then stained with a 1:1000 dilution of Hoechst 33342 (Thermo Scientific, 62249) for 15 minutes, then washed twice with 1x PBS, and mounted on glass slides using ProLong Diamond antifade mountant (Invitrogen, P36961). Slides were sealed using clear nail polish, allowed to dry, and stored at 4°C.

Slides were imaged on either an Olympus FV1200 Laser Scanning confocal microscope or a DeltaVision 2 TIRF microscope. All partition and size analyses were performed using images from the Olympus FV1200 confocal microscope. Costains of transfected human cells were imaged on either microscope. Coassembly experiments were imaged on the DeltaVision 2 TIRF microscope. Zebrafish cells derived from injected embryos were imaged on the DeltaVision 2 TIRF microscope. The same set of exposure conditions (one exposure per channel) was used across all slides within the same experiment. Maximum projections across the z-axis of acquired images were made using Fiji (https://imagej.net/software/fiji/) and used for any further analyses.

### Partition and size analysis

Maximum projection images were analyzed to calculate average nuclear intensity, the free energy of transfer (ΔG^transfer^), and dense phase size of GFP-tagged proteins using custom python code (found on zenodo at: https://doi.org/10.5281/zenodo.7236056). Briefly, nuclei were segmented in the images by applying gaussian blur, followed by thresholding at the mean intensity in the Hoecsht 33342 channel.

The average nuclear intensity for each nucleus was defined as the average nuclear GFP signal following background correction, and was used as a proxy for nuclear concentration of the GFP-tagged protein. Background signal was defined as the 75th percentile of GFP signal in nuclei of an untransfected sample. This background value was subtracted from the GFP intensity of all images of transfected samples within the set of images taken with identical settings. GFP signal beyond this point refers to the background corrected GFP signal.

To calculate ΔG^transfer^, which is a measurement of the energy of partitioning into the dense phase, the dense and dilute phase were determined by segmenting the GFP channel using Otsu thresholding. Nuclei in which the Otsu threshold was below an intensity value of 100 were considered untransfected cells and excluded from analysis. This threshold was determined by manually checking that untransfected cells were excluded based on the value of the threshold. Any nuclei with overexposed pixels in the GFP channel were also excluded from analysis. For partition analysis, dense phase concentrations were defined as the 90th percentile GFP intensity within the segmented region, and dilute phase concentrations were defined as the median GFP intensity outside of the segmented region. ΔG^transfer^ was then calculated using the following equation:

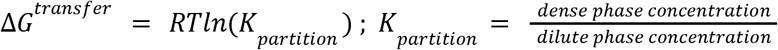

where R is the gas constant 1.987 × 10^−3^ kcal K^-1^ mol^-1^ and T = 310.15 K. Dense phase size was defined as the number of pixels in the dense phase as segmented above.

In some nuclei, the low intensity of dilute phase after background correction resulted in noisy partition coefficient calculations, so we used a minimum dilute phase intensity cutoff intensity of 6 to mitigate this. ΔG^transfer^ was not calculated for nuclei which did not pass this cutoff, as this calculation requires a dilute phase signal. However the dense phase size calculation does not require a dilute phase signal, and as such this calculation was still performed on these nuclei. As a consequence, the number of nuclei used for the ΔG^transfer^ calculation for some experiments is smaller than the number of nuclei used for the dense phase size calculation.

Fits for partition analyses were performed using the regplot function in seaborn, with “logx=True” for ΔG^transfer^ plots, and “logx=False” for dense phase size plots. For all fits, 95% confidence intervals are shown.

### Transmission electron microscopy (TEM)

Cells were seeded in 6-well cell culture plates. Upon collection, media was removed from the plates and immediately fixed using 2.5% Glutaraldehyde 2.5% Paraformaldehyde in 0.1 M sodium cacodylate buffer at pH 7.4 (Electron Microscopy Sciences, 15949) for at least 1 hour at room temperature before embedding.

For embedding, fixative was removed and the cells were washed twice in 0.1 M Sodium Cacodylate buffer, pH 7.4. Then, the cells were incubated in 1% Osmium tetroxide (OsO_4_)/1.5% Potassium ferrocyanide (KFeCN_6_) for 30 min, washed twice in water, and once in 50mM Maleate buffer, pH 5.5 (MB). The plate was then incubated in 1% uranyl acetate in MB for 30 min followed by one wash in MB and two washes in water and subsequent dehydration in grades of alcohol (5min each; 50%, 70%, 95%, 2x 100%). Following the washes, cells were removed from the dish in propyleneoxide, pelleted at 3000 rpm for 3 min and incubated overnight in a 1:1 mixture of propyleneoxide and TAAB Epon (TAAB Laboratories Equipment Ltd, https://taab.co.uk). The following day the samples were embedded in TAAB Epon and polymerized at 60°C for 48 hrs.

Ultrathin sections (about 60-80nm) were then cut on a Reichert Ultracut-S microtome, picked up on copper grids, stained with lead citrate and examined in a JEOL 1200EX Transmission electron microscope at 4000X. Images were recorded with an AMT 2k CCD camera.

### Imaging and quantification of nascent transcription in nucleoli

Nascent transcription in nucleoli was measured by incorporation of 5-ethynyl uridine (5-EU) in cells using a Click-iT™ RNA Alexa Fluor™ 594 Imaging Kit (Fisher Scientific, C10330), according to the manufacturer’s recommended protocol except where noted in the following. Briefly, cells were plated on fibronectin-coated glass coverslips in 24-well plates, as described above. On the day of collection, cells were incubated in 1 mM 5-EU for 3 hours at 37°C. After 5-EU incorporation, cells were washed once with 1x PBS, and then fixed in 4% paraformaldehyde for 15 minutes at room temperature. Permeabilization, and Click reaction to covalently attach Alexa 594, and washes were all done as recommended by the kit. After washing the Alexa 594 Click reaction, but before Hoechst 33342 nuclear staining, cells were stained for nucleolar marker MPP10 using the immunofluorescence and imaging methods as described above.

To quantify nascent transcription in cells, nuclei of cells were segmented, followed by segmentation of nucleoli, and measurement of 5-EU Alexa 594 signal using custom python code (found on zenodo at: https://doi.org/10.5281/zenodo.7236056). Using max projection images, nuclei were segmented using the Hoechst 33342 channel as previously described above, and nucleoli were segmented using the MPP10 channel by performing Otsu thresholding on nuclear MPP10 signal. Then, for each nucleus, the amount of nascent transcription in nucleoli was calculated as the average 5-EU intensity within MPP10 segmented region/s. Background correction was performed by subtracting the mean nucleolar 5-EU intensity of a sample that was not treated with 5-EU. These data were then plotted to compare different cell lines and conditions. A two-tailed t-test with no assumption of equal variance was used to test statistical significance.

### In vitro transcription of RNA

To generate the RNA for zebrafish microinjections in Fig. S10B, a construct encoding meGFP-TCOF1 (RP133) was amplified using Q5 DNA polymerase (New England BioLabs, M0491L) reaction mix with primers RPO378 (GCGTAATACGACTCACTATAGGGAGACCCAA) and RPO379 (GTGGATCCGAGCTCGGTACCAA). Products were then run on a 1% agarose gel and the gel was extracted using NucleoSpin (Macherey-Nagel, 740588.50) clean up kit according to manufacturer specifications.

For Fig. 4C, RNA was generated from digesting 2μg of RP133 and RP168 with BamHI-HF (New England BioLabs, R3136S) for 1hour. The product was then purified from 1% agarose gel and extracted using the NucleoSpin clean up kit (Macherey-Nagel, 740588.50).

500 ng and 300 ng of PCR DNA and digested linearized DNA, respectively, were used to set up an overnight (16 hours) IVT reaction using mMESSAGE mMACHINE T7 ULTRA Transcription Kit (Invitrogen, AM1345). The overnight product was then treated with TURBO DNase, from the same mMESSAGE mMACHINE Kit, for 30min. After the DNAse treatment the samples were poly-A tailed using the tailing reaction provided in the mMESSAGE mMACHINE Kit. 2μL of the DNA was removed prior to adding E-PAP enzyme, to be run at a later time point on RNA agarose gel to confirm band size and act as control for post poly-A tailing reaction. Samples were then purified using Zymo RNA Clean and Concentrator kit (Zymo Research, R1016).

### TCOF1 ortholog search approach and definition

To search for the existence of orthologs of TCOF1 in other species, we used the BLAST webtool from UniProt (https://www.uniprot.org/blast) and NCBI (https://blast.ncbi.nlm.nih.gov/Blast.cgi), allowing for maximal returned hits and no filtering based on low complexity sequences. Because of the low complexity of human TCOF1, orthologs of TCOF1 were determined based on manual assessment of the sequence rather than by alignment coverage. Species which contained TCOF1 were defined as species where a sequence from the BLAST results was similar to human TCOF1 based on manual dotplot analysis of the LCR relationships within each given protein sequence. In particular, TCOF1 orthologs were defined based on the presence of repeat domains containing multiple S/E-rich LCRs and a C-terminal region containing K-rich LCR(s).

### Zebrafish husbandry

The zebrafish were handled and maintained according to the vertebrate animal protocol. AB wild-type strains were used.

### Injection of zebrafish embryos

Zebrafish were crossed by placing a male and female fish in a shared container, but kept physically separate until combined the next morning. Embryos were collected for injection 20 minutes after males and females were combined for fertilization. Injection mix consisting of 0.28 mM *in vitro* transcribed RNA and 0.07% Phenol Red solution (Sigma, P0290) was prepared and loaded into needles pulled from glass capillary tubing (64-0766). Embryos were injected 1 nL of injection mix using a pico-liter injector (Warner Instruments, PLI90A). Injected embryos and uninjected controls were then incubated in fish water at 28°C until collection at 24 hours post fertilization (hpf).

### Whole-mount imaging of zebrafish embryos

Zebrafish embryos at 24 hpf were collected, dechorionated using ethanol-sterilized forceps, and moved to fresh fish water. Using a transfer pipette, individual embryos were anesthetized by transfer into 1X Tricane solution, prepared by diluting 25 X Tricane solution (pH 7 solution of 4mg/mL Tricane-S (Syndel) and 20 mM TRIS) to 1 X in fish water. Fish were subsequently transferred into 1.5% low-melt agarose (Lonza, 50100), and transferred to a clear dish suspended in low-melt agarose for imaging. Bright-field and GFP fluorescence images were taken on a Leica M205 FCA fluorescence stereo microscope. Images were taken with the same settings for all embryos, at 5.5 X magnification.

### Deriving cells from zebrafish embryos

Cells for immunofluorescence were derived from 24 hpf embryos grown at 28 C. Embryos expressing high GFP signal were selected prior to dechorionation. Embryos were dechorionated, using ethanol-sterilized forceps for immunofluorescence or pronase (1mg/mL) for electron microscopy, and then transferred to 1x PenStrep Glutamine (Gibco, 10378016) in 1x PBS (PBS+PSG) for 30 minutes. Once the embryos were dechorionated sterile tubes and pipettes tips were used to minimize any possible contamination to the embryos/cells. Embryos were then transferred to new tubes with fresh PBS+PSG. Yolk-sacs were removed through pipetting several times with P200 pipette. Embryos were then spun at 1200 x g for 2 min at room temperature and the supernatant was then discarded. The embryos were then incubated in 300μL of prewarmed .25% Trypsin (Gibco, 25200072) for 30 minutes at 1000 rpm on a thermomixer (Eppendorf, Thermomixer R). The embryos were then pipetted with P200 to further resuspend the embryos into single cell suspension and then were spun at 1200g for 4 minutes at room temperature. Supernatant was disposed of and the embryos were serially washed with sterile PBS (Genesee Scientific, 25-508). Cells were then resuspended in 400μL of DMEM, supplied with 15% FBS (Gemini Bio, 100-106), 1X PSG (Gibco, 10378016) and 1 mg/mL gentamicin (Gibco, 15750060) in 24-well cell cultures plates (Genesee Scientific, 25-107) with coverslips (Fisher Scientific, 1254580) that were coated with 3μg/mL fibronectin in PBS for at least 30 minutes prior to plating cells. Cells were allowed to attach overnight at 28 C supplied with 5% carbon dioxide prior to immunofluorescence staining.

**Fig. S1.**
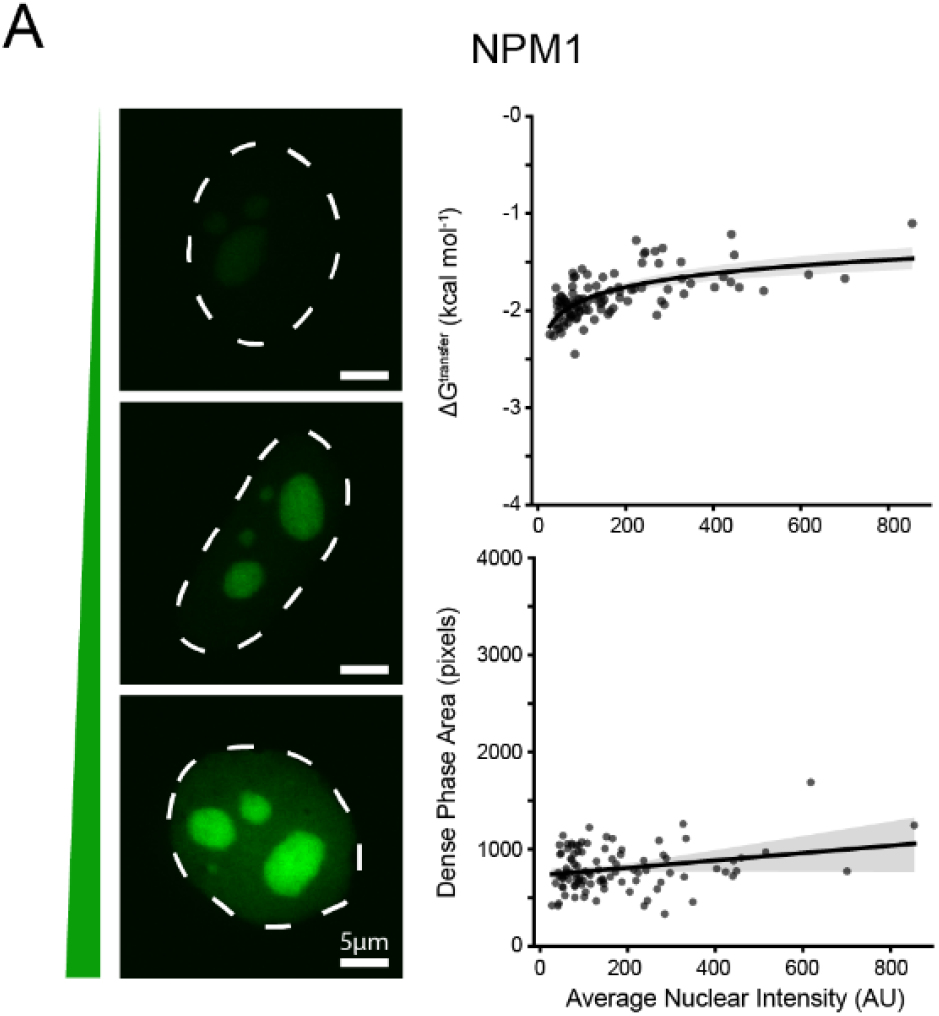
NPM1 displays multi-component assembly in cells. Quantification of ΔG^transfer^ and dense phase size vs. average nuclear intensity for NPM1. Images of cells with different expression levels are shown on the left. Nuclei are outlined with a dotted line based on Hoechst 33342 signal (not shown). Scale bars are 5 μm. N = 100 nuclei ΔG^transfer^ measurement, and N = 100 nuclei for dense phase area measurement. Logarithmic and linear fits with corresponding 95% confidence intervals are shown.

**Fig. S2.**
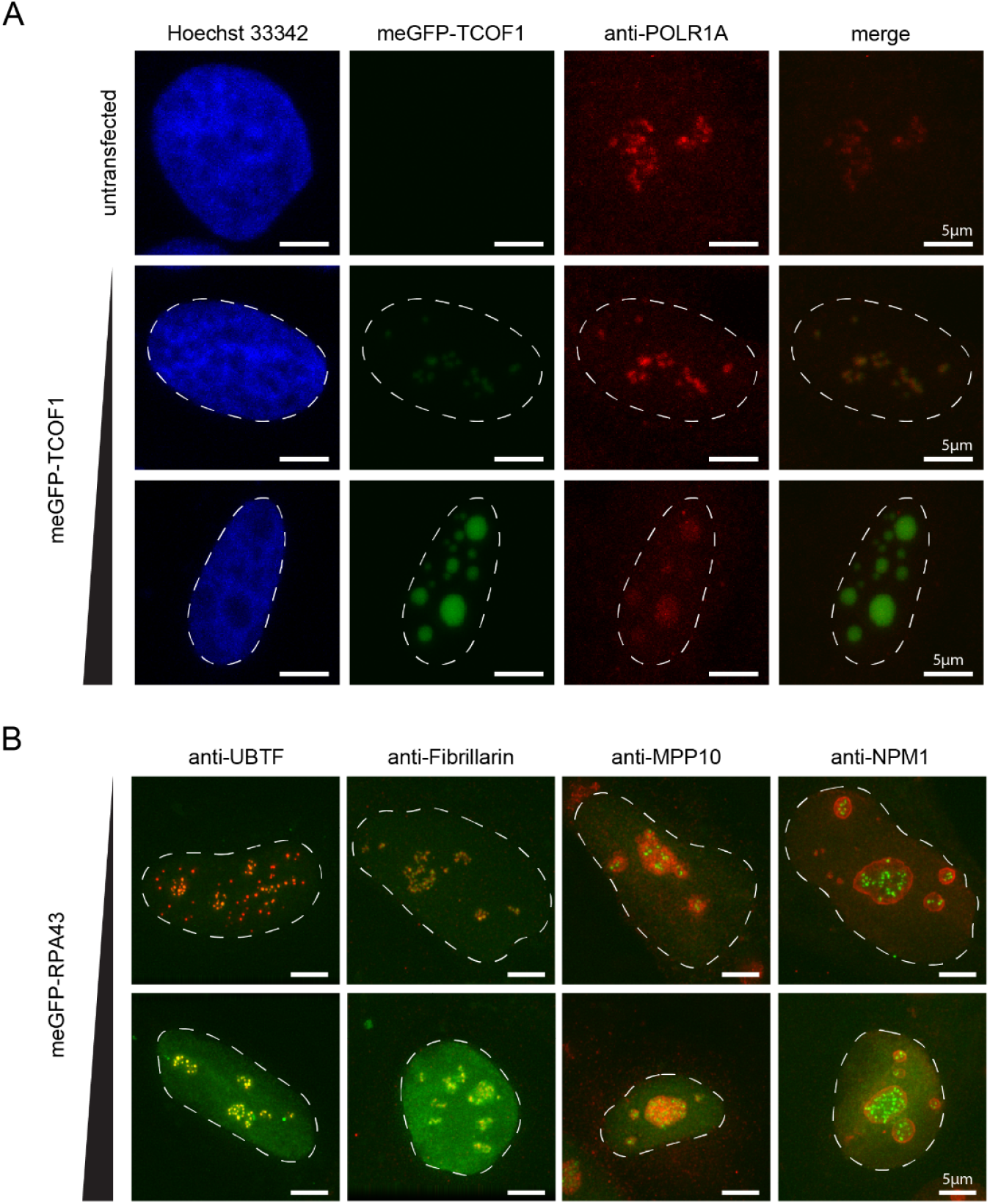
Supplementary Data for FC expansion. A) Immunofluorescence of the fibrillar center marker POLR1A across varying levels of meGFP-TCOF1 expression. Nuclei are outlined with a dotted line based on Hoechst 33342 staining. Scale bars are 5 μm. B) Immunofluorescence of endogenous nucleolar subcompartment markers across varying levels of meGFP-RPA43 expression. Nuclei are outlined with a dotted line based on Hoechst 33342 staining (not shown). Scale bars are 5 μm.

**Fig. S3.**
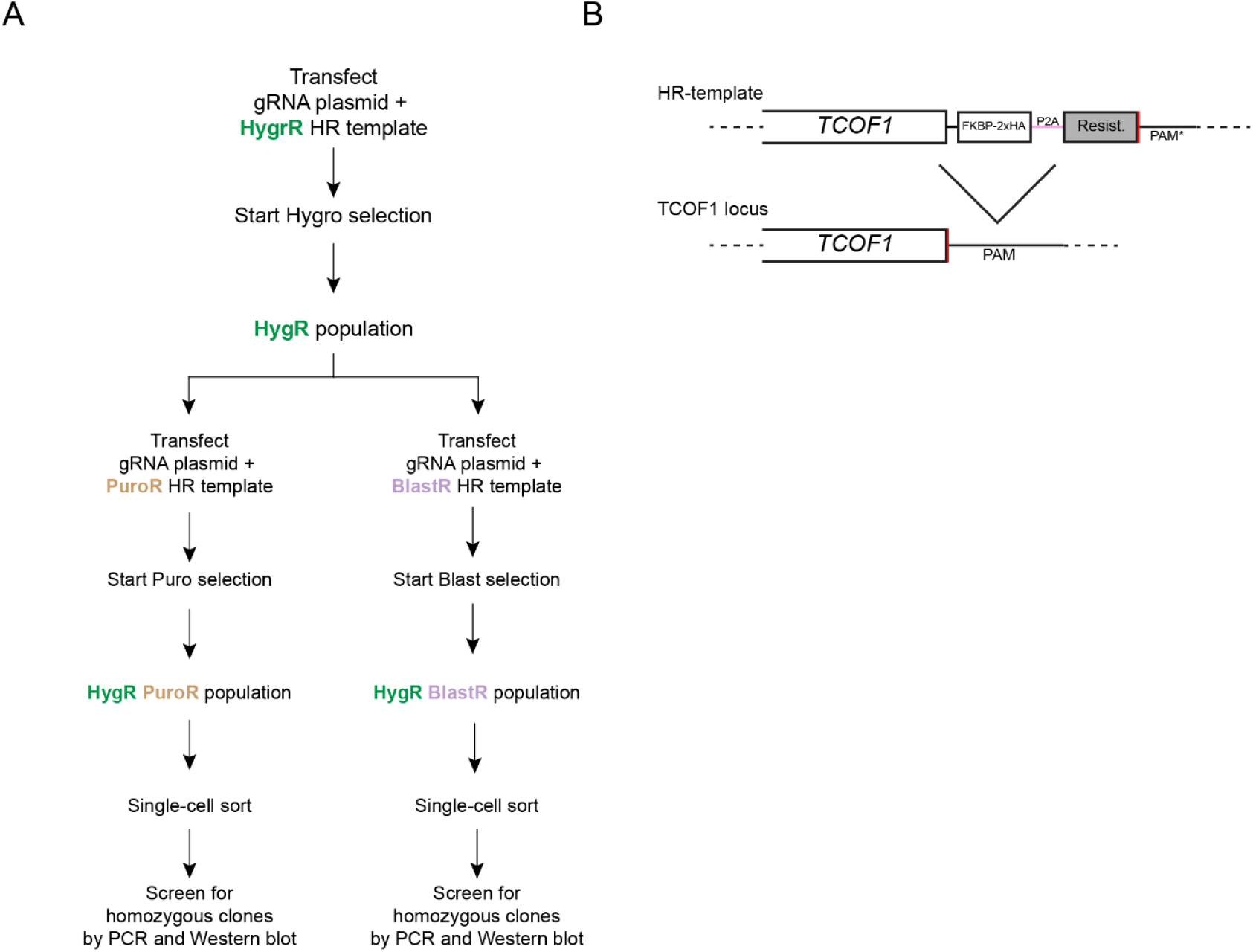
Generation of TCOF1 degron cell lines. A) Scheme describing generation of TCOF1 degron lines. See Methods for details. B) Schematic of homologous recombination (HR) templates used for generation of TCOF1 lines.

**Fig. S4.**
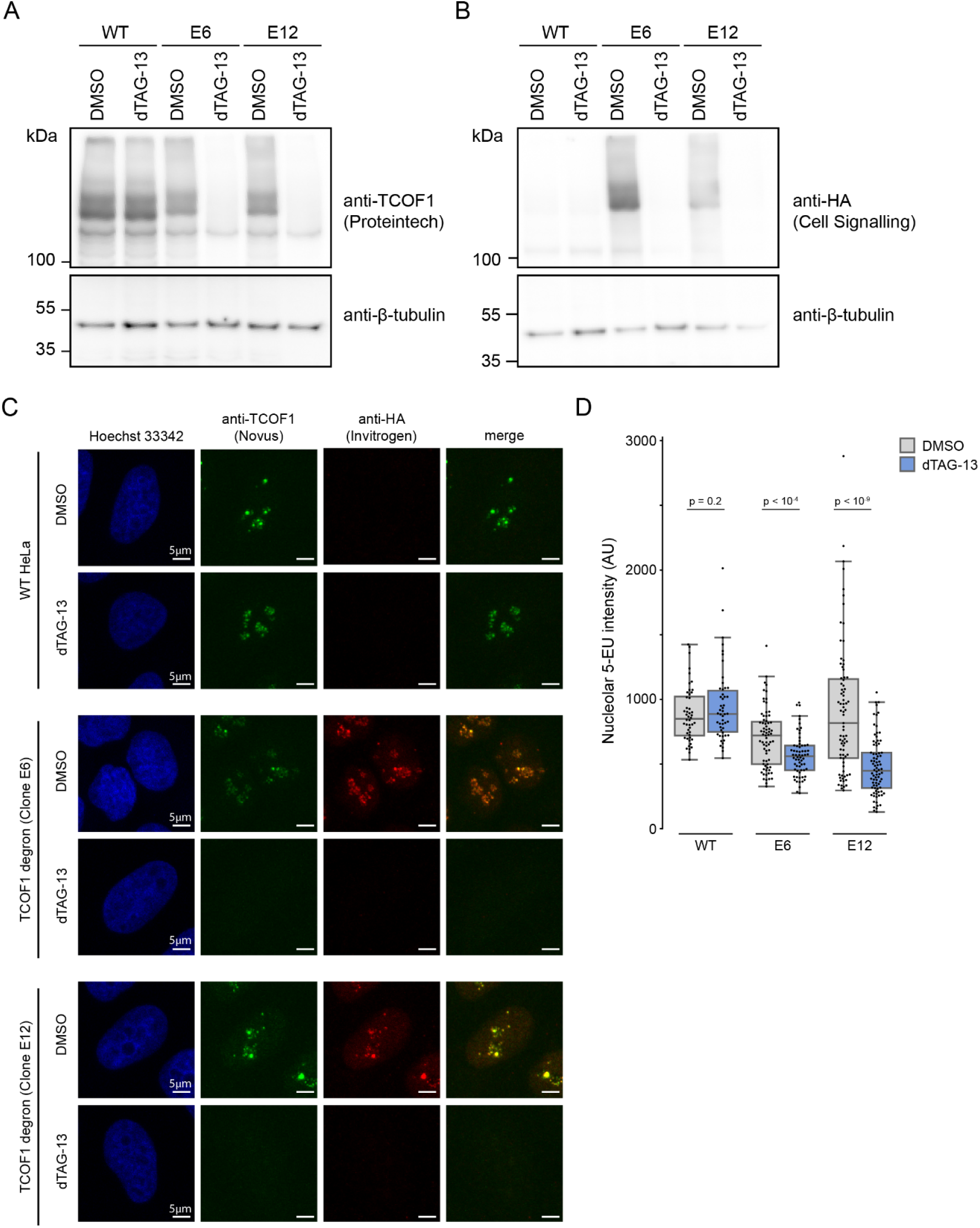
Validation of TCOF1 degron cell lines. A) Western blot of DMSO and dTAG-13 treated WT HeLa cells and TCOF1 degron cells using TCOF1 antibody (Proteintech, 11003-1-AP) and β-tubulin antibody as loading control. B) Western blot of DMSO and dTAG-13 treated WT HeLa cells and TCOF1 degron cells using HA antibody (Cell Signaling (C29F4), 3724S) and β-tubulin antibody as loading control. C) Immunofluorescence images of DMSO and dTAG-13 treated WT Hela cells and TCOF1 degron cells stained using TCOF1 antibody (Novus Biologicals, NBP1-86909) and HA antibody (Invitrogen, 26183). Scale bars are 5 μm. D) Quantification of nascent nucleolar RNA in WT HeLa cells and HeLa degron lines after treatment with DMSO or dTAG-13 for 4-days. A two-tailed t-test with no assumption of equal variance was used to test statistical significance between DMSO and dTAG-13 treated samples for each cell line, and p-values are indicated. N = 48 nuclei for WT DMSO, N = 50 nuclei for WT dTAG, N = 71 nuclei for E6 DMSO, N = 64 nuclei for E6 dTAG, N = 79 nuclei for E12 DMSO, N = 82 nuclei for E12 dTAG.

**Fig. S5.**
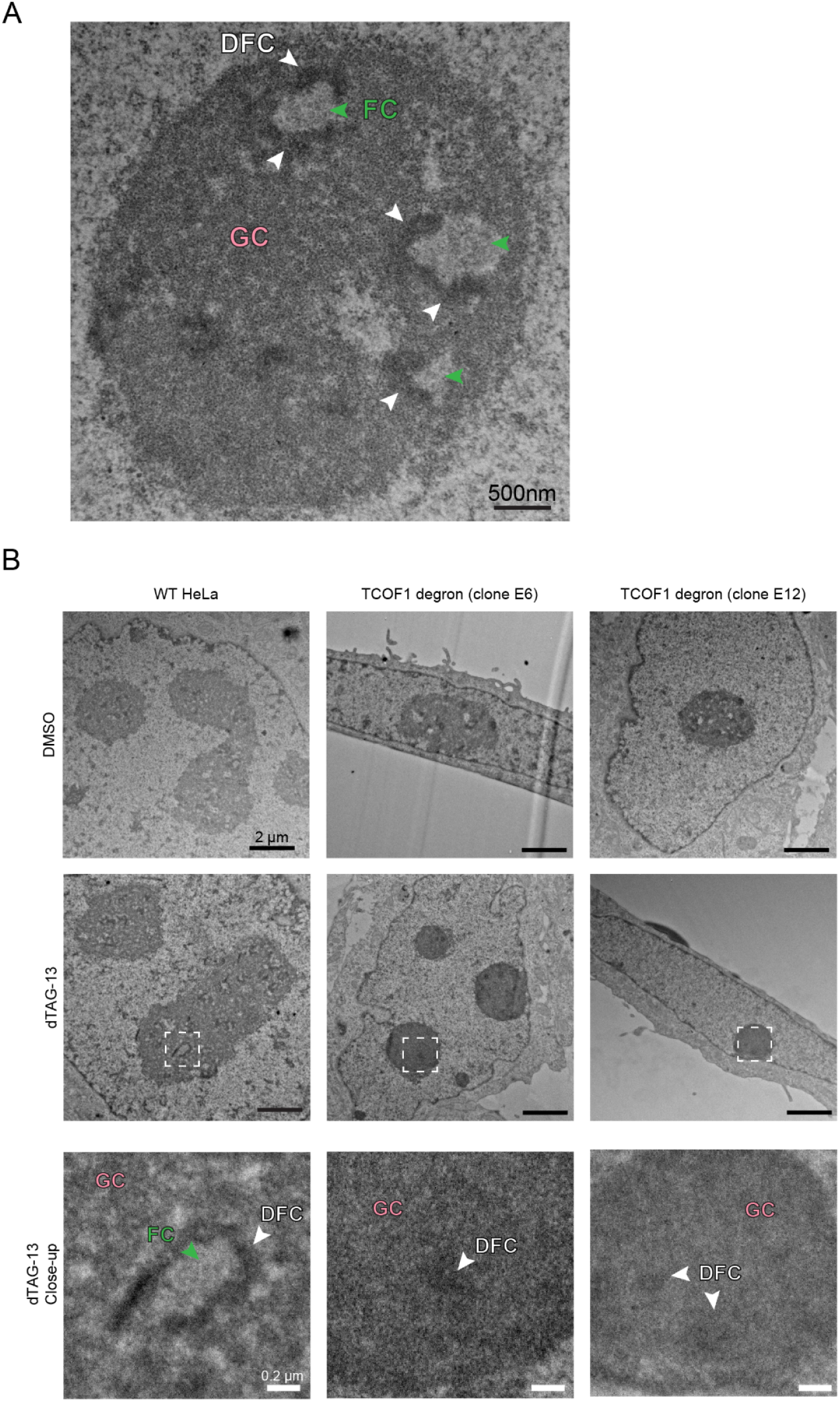
Supplementary data for EM of TCOF1 degron lines. A) Transmission electron microscopy image of a nucleolus with the GC, DFC, and FC labeled with colored arrowheads. B) Transmission electron microscopy of WT HeLa cells and two different clones of TCOF1 degron HeLa cell lines after 4-days of DMSO or dTAG-13 treatment. Bottom row is a close-up view of region indicated with white box in images of dTAG-13 treated cells in the second row, with regions corresponding to FC, DFC, and GC indicated. Scale bars are 2 μm for the top two rows, and 0.2 μm for the bottom row.

**Fig. S6.**
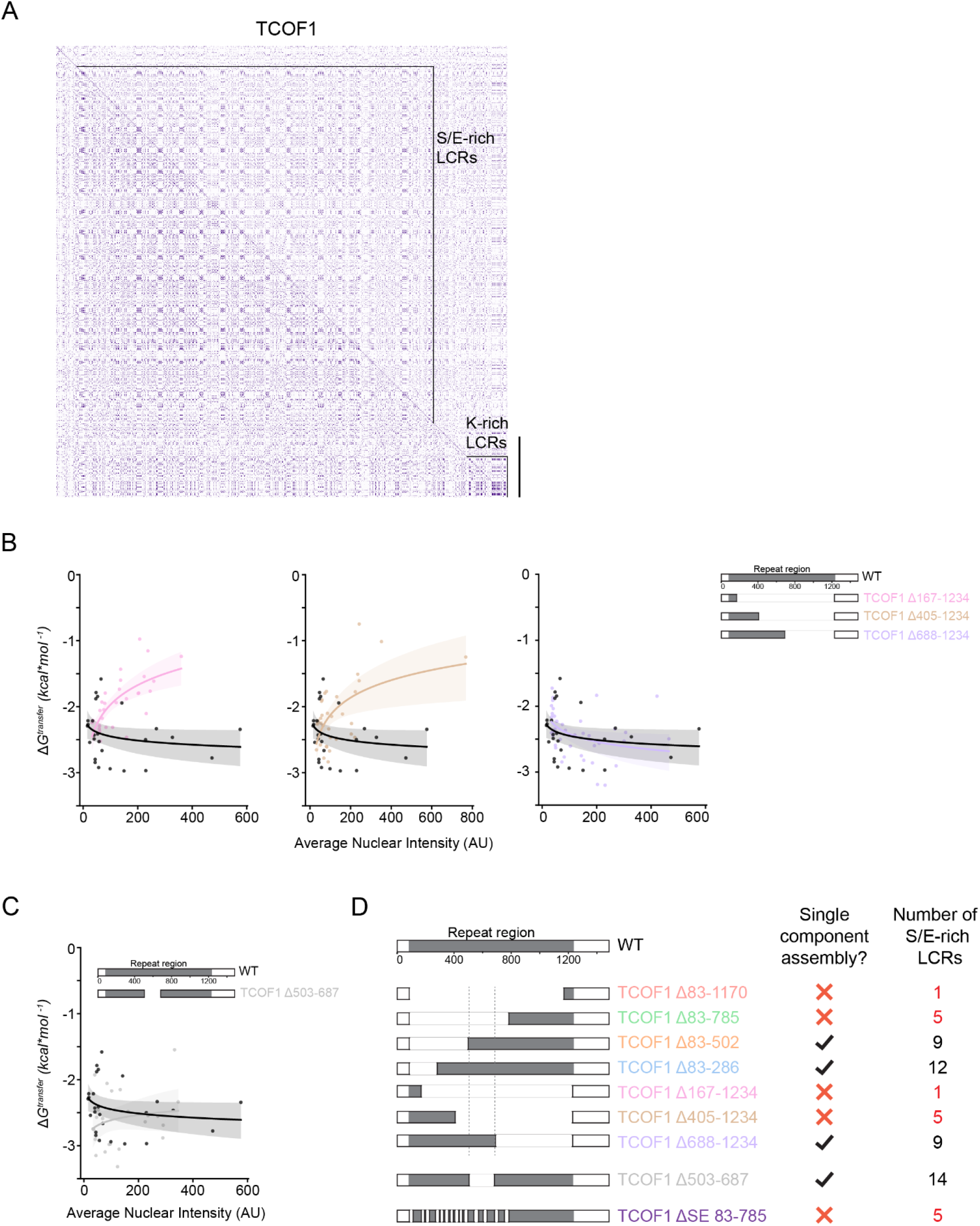
Supplementary data for partition analysis of TCOF1 mutants. A) Dotplot of human TCOF1. B) Quantification of ΔG^transfer^ vs. average nuclear intensity for WT TCOF1 and TCOF1 mutants with internal deletions in the central repeat region. Data for WT TCOF1 is shown on each plot for clarity when comparing between WT and mutants. N = 26 nuclei for WT TCOF1, N = 29 nuclei for TCOF1 Δ167-1234, N = 34 nuclei for TCOF1 Δ405-1234, N = 40 nuclei for TCOF1 Δ688-1234. Logarithmic fits and corresponding 95% confidence intervals are shown. C) Quantification of ΔG^transfer^ vs. average nuclear intensity for WT TCOF1 and a TCOF1 mutant with an internal deletion of a region which was present in all mutants which displayed single component assembly in Figure 3A and Figure S6A. Data for WT TCOF1 is shown on each plot for clarity. N = 26 nuclei for WT TCOF1, N = 24 nuclei for TCOF1 Δ503-687. Logarithmic fits and corresponding 95% confidence intervals are shown. D) Schematic illustrating all TCOF1 mutants tested, their assembly behavior, and the number of S/E-rich LCRs they possess.

**Fig. S7.**
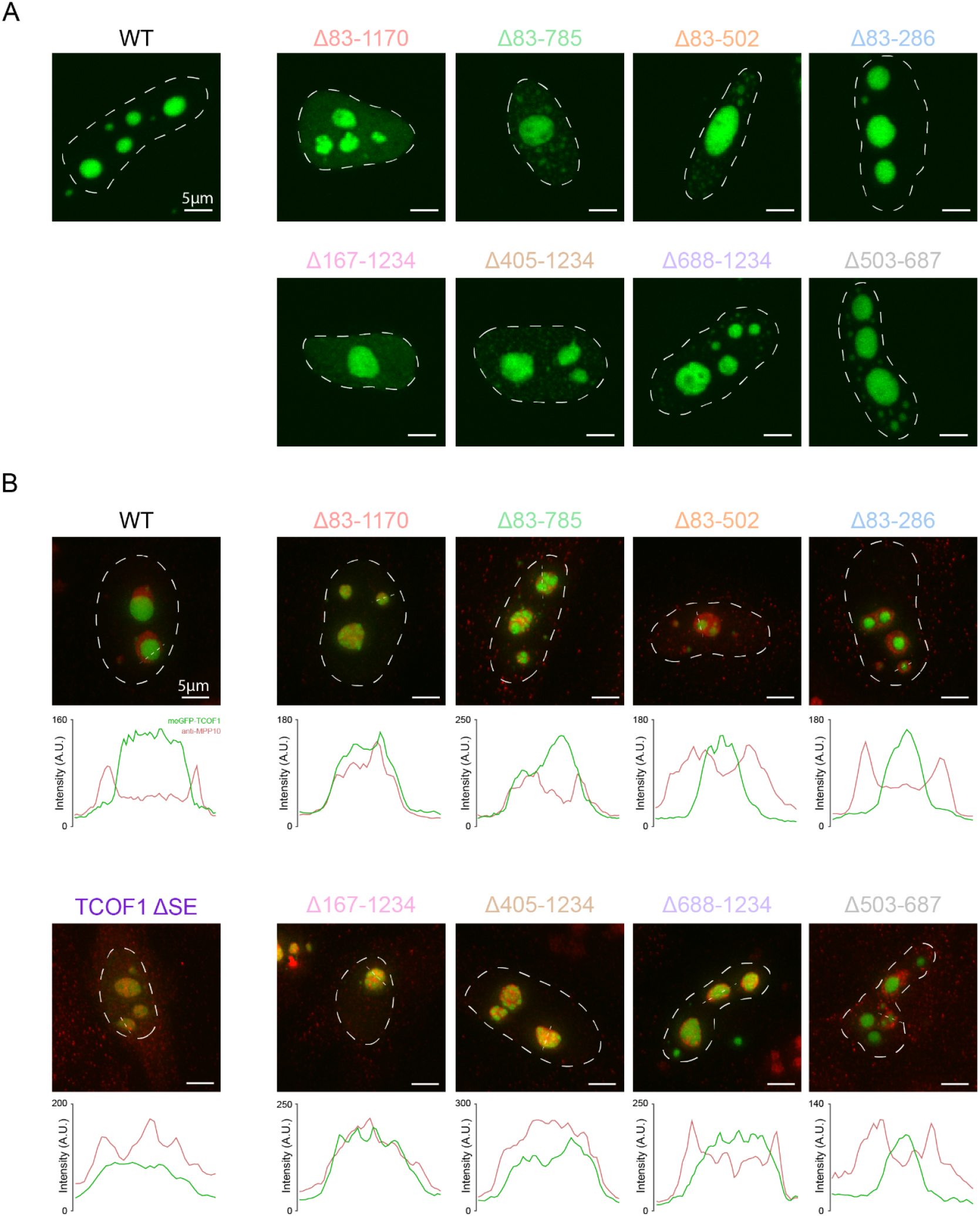
Supplementary data for imaging of TCOF1 mutants. A) Representative fluorescence images of cells expressing WT TCOF1 and different TCOF1 mutants as labeled. All images were taken at the same exposure. Scale bars represent 5 μm. B) Representative fluorescence images of localization of nucleolar marker MPP10 with meGFP-WT TCOF1 or different meGFP-TCOF1 mutants as labeled. Nuclei are outlined with a dotted line based on Hoechst 33342 signal (not shown). Scale bars represent 5 μm. Line profile plots are included over the region indicated by the thin dotted line for meGFP (green) and MPP10 channels (red).

**Fig. S8.**
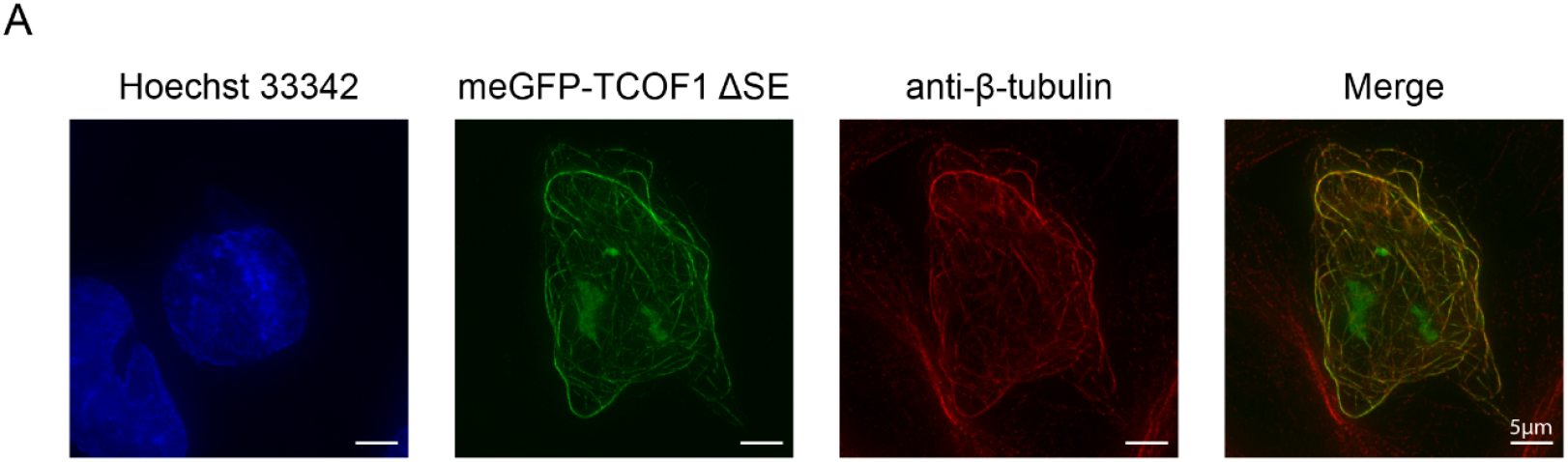
Supplementary data for TCOF1 ΔSE mutant. A) Immunofluorescence of HeLa cells transfected with meGFP-tagged TCOF1 ΔSE and stained for β-tubulin. Scale bars are 5 μm.

**Fig. S9.**
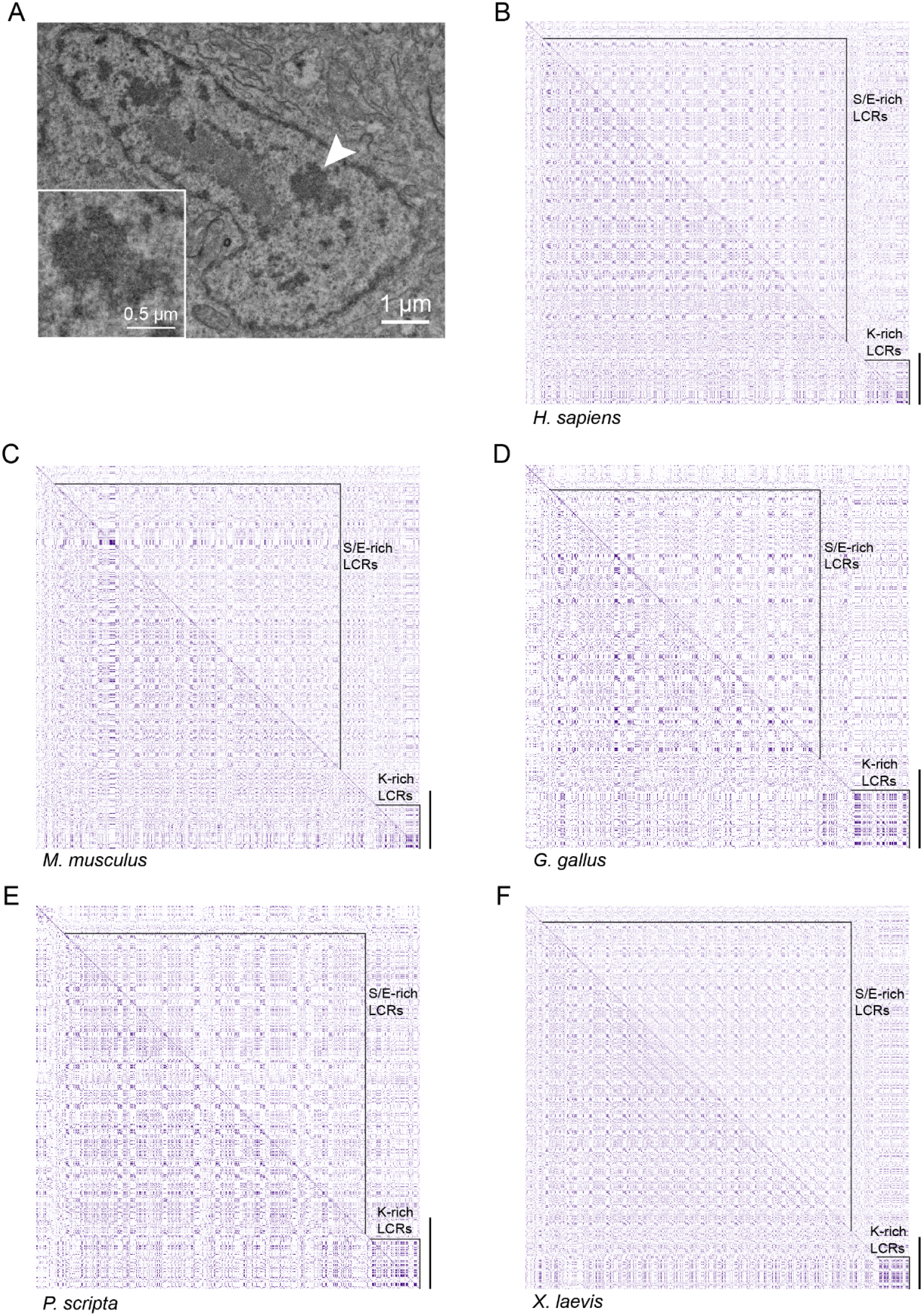
Supplementary data for analysis of FC/TCOF1 over evolution. A) Transmission electron microscopy of cells derived from zebrafish embryos. Scale bar is 1 μm. Inset shows close-up view of region indicated by arrowhead. Inset scale bar is 0.5 μm. B) Dotplot of human TCOF1. C-F) Dotplots of TCOF1 orthologs in *M. musculus, G. gallus, P. scripta*, and *X. laevis*, respectively. Overlays indicate S/E-rich LCRs and K-rich LCRs as annotated. Scale bar represents 200 amino acids in length.

**Fig. S10.**
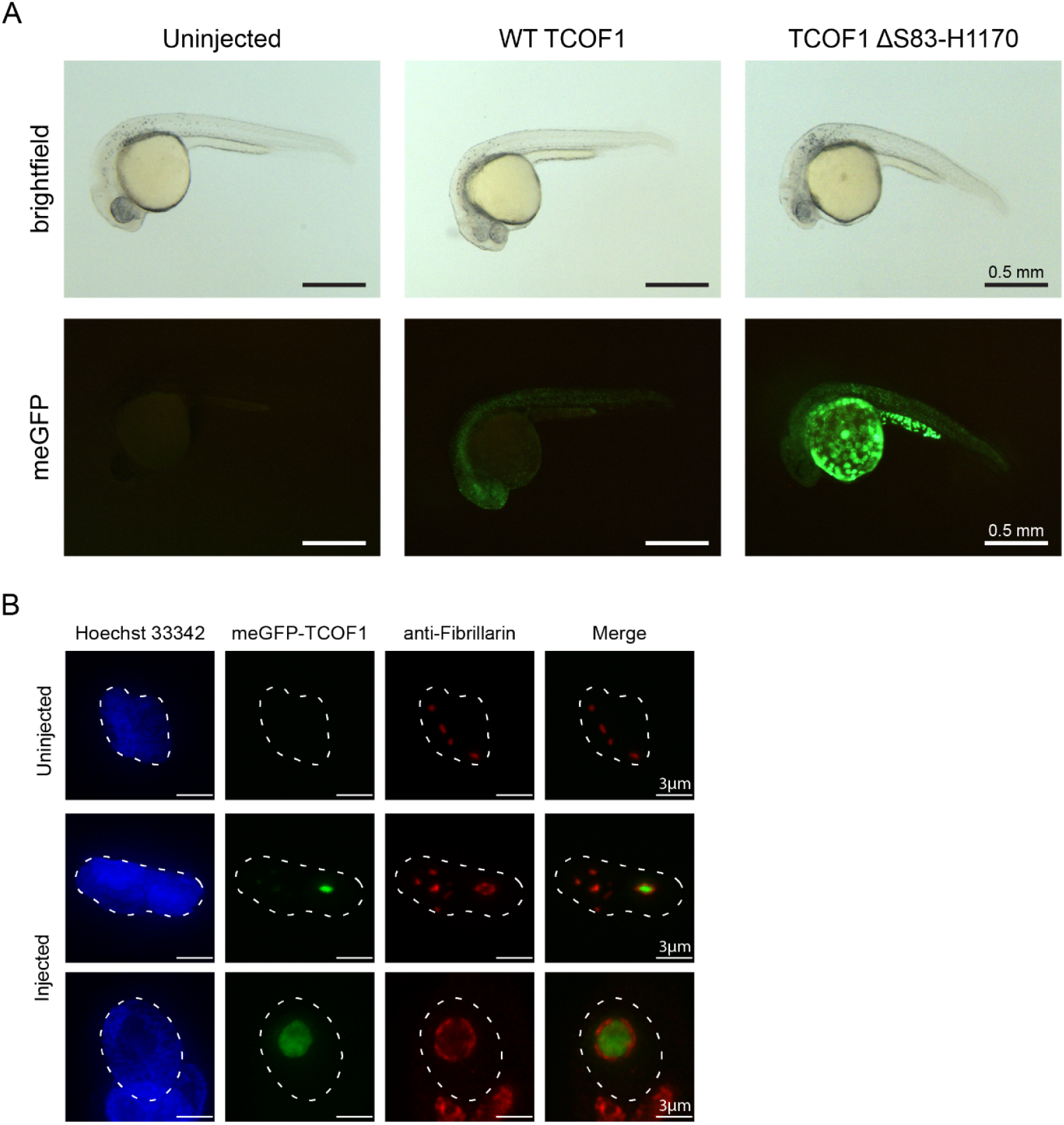
Supplementary data for zebrafish injection. A) Whole-mount zebrafish embryo images of an uninjected control and embryos injected with WT TCOF1 and TCOF1 ΔS83-H1170. B) Fluorescence images of uninjected control and embryos injected with WT TCOF1 at low and high expression levels. Nuclei are outlined with a dotted line based on Hoechst 33342 signal. Scale bars are 3 μm.

**Table S1.** Table summarizing presence of TCOF1 orthologs and nucleolar structure across species

| Species | TCOF1 ortholog | Tripartite/bipartite | Reference |
| --- | --- | --- | --- |
| *H. Sapiens* | TCOF_HUMAN (Uniprot) | Tripartite | Cheutin *et al.* 2002. Journal of Cell Science; This study (Fig. 2C) |
| *M. musculus* | TCOF_MOUSE (Uniprot) | Tripartite | Mirre and Knibiehler 1982, Journal of Cell Science |
| *G. gallus* | F1NRG3 (Uniprot) | Tripartite | Macera and Bloom. 1981, The journal of heredity |
| *P. scripta* | XP_034635347.1 (NCBI) | Bipartite | Lamaye *et al.* 2011. Journal of Structural Biology |
| *X. laevis* | Q5G8Z4 (Uniprot) | Tripartite | Mais and Scheer. 2001, Journal of Cell science |
| *D. rerio* | Not found | Bipartite | Selman *et al.* 1993, Journal of Morphology; this study (Fig. S9A) |
| *D. melanogaster* | Not found | Bipartite | Knibiehler, Mirre, and Rosset 1982, Journal of Cell Science |
| *S. cerevisiae* | Not found | Bipartite | Lýger-Silvestre *et al.* 1999. Chromosoma |

